# Hive position matters: spatial layout and hive colour impact forager traffic and drifting in *Melipona quadrifasciata* bees

**DOI:** 10.64898/2026.08.01.742195

**Authors:** Ana Paula Cipriano, Cristiano Menezes, Milly Munro, Emily Bell, Christoph Grüter

## Abstract

Stingless bees are key pollinators of native and cultivated plants, and their management mostly for honey and hive commercialisation has become an increasingly popular practice in Brazil. Hives are commonly kept close together and visually similar in meliponaries, a practice that favours drifting, i.e., the return of foragers to a foreign nest. Distance between hives and layout have been identified as drivers of drifting in the Western honeybee, but evidence for stingless bees, the largest group of social bees, remains scarce. Additionally, stingless beekeepers frequently associate drifting with weakened colonies, uneven honey production, worker fighting, and colony population imbalances. We investigated the effects of hive layout on foraging traffic and drifting in the stingless bee *Melipona quadrifasciata*. We compared forager traffic across six layouts varying in distance, colour and entrance orientation. Then, using RFID tags, we tracked drifting across three layouts with the same inter-hive distance but different designs: visually similar hives, hives with opposite entrance orientation, and hives painted with distinct colours. Forager traffic was lowest in the visually similar layout during the first two weeks of monitoring, which is consistent with greater disorientation in the absence of clear visual cues. Hive position was a key driver of drifting behaviour across all layouts, with bees from edge hives drifting considerably less than those from inner hives. Drifting decreased considerably in hives with different colours (37.6%) and hives with entrances in opposite directions (48.1%), compared to similarly looking hives with entrances in the same direction (59.5%). Therefore, our findings highlight practical low-effort strategies for stingless beekeepers to reduce forager loss, pathogen spread, and productivity losses.

## Introduction

Stingless bees (Meliponini) play key roles as pollinators of native and cultivated plants (Bueno et al., 2023; Giannini et al., 2015; Roubik, 2018), and are distributed in tropical and subtropical ecosystems in the Americas, Africa, Asia and Oceania, with more than 600 species described (Engel et al., 2023; Grüter, 2020; Lepeco et al., 2024; Quezada-Euán, 2018). In many of these regions, stingless beekeeping (i.e., meliponiculture) has been practised by diverse communities (Cortopassi-Laurino et al., 2006), including the Mayans (Quezada-Euán, 2018), the Kayapó in Brazil, Aboriginal communities in Australia and communities in Tanzania and Angola (Cortopassi-Laurino et al., 2006), but meliponiculture used to be a small-scale and largely informal activity (Jaffé et al., 2015). However, in recent years, keeping stingless bees for the commercialisation of honey and hives has become an increasingly popular and professional practice in different regions of Brazil, where bee products from multiple species now constitute a profitable activity (Imperatriz-Fonseca et al., 2017; Jaffé et al., 2015; Leão et al., 2025; Pereira et al., 2017). Despite its antiquity, importance, and growing popularity, meliponiculture still requires a better understanding of best practices to guarantee sustainable hive productivity, that is, by also prioritising bee health.

In natural conditions, stingless bee nest density varies from as low as 0.014 nests/ha to 1.1 nests/ha (Silva and Ramalho, 2016). However, in managed areas (i.e., meliponaries), several hives are commonly kept close together and designed to be visually similar (Leão et al., 2025). This hive arrangement favours drifting, which occurs when a bee returns to the wrong hive after foraging (Bordier et al., 2017; Free, 1958; Oliveira et al., 2021; Stephens et al., 2017). The distance between hives and the layout are important factors driving drifting rates in the Western honeybee (*Apis mellifera*) (Jay, 1966; Pfeiffer and Crailsheim, 1998; Seeley and Smith, 2015), the bee species most studied in relation to drifting. In this species, positioning hives 1m apart in a row with no visual distinction can result in drifting rates of 25% up to 46% (Dynes et al., 2019; Seeley and Smith, 2015). In contrast, in apiaries with dispersed hives 30m apart, or hives 10m apart, but painted with different colours and positioned in a circle at different heights, the drifting percentages decrease to 0% and 7.5%, respectively (Dynes et al., 2019; Seeley and Smith, 2015). Maintaining *A. mellifera* hives in crowded apiaries (i.e., a large number of hives positioned 1m apart) has also been associated with higher parasite levels (Dynes et al., 2019; Seeley and Smith, 2015), reduction in honey production (Dynes et al., 2019), and, indirectly, colony death (Dynes et al., 2019; Seeley and Smith, 2015). Additionally, in a simulated environment, bee drifting patterns were an emergent property of navigational mistakes, with inter-hive distance substantially reducing drifting rates, and hive position affecting bee population and energy accumulation (Cipriano et al., 2026). Both honeybees and stingless bees tend to drift more from central hives and visit their closest neighbour at a high rate (Dynes et al., 2019; Forfert et al., 2015; Oliveira et al., 2021; Pfeiffer and Crailsheim, 1998; Stephens et al., 2017), which may cause an imbalance in hive population and food stores as well as a higher disease risk for certain hives within the apiary.

A better understanding of drifting in social bees can improve management practices that favour colony health and honey production, but evidence for stingless bees, the largest group of social bees, remains scarce. Data on *Tetragonula carbonaria* and *Melipona fasciculata* bees indicate high drifting rates in managed hives, with up to 42% and 65%, respectively (Oliveira et al., 2021; Stephens et al., 2017), compared to 0% in natural nests (Stephens et al., 2017). This contrast highlights a clear influence of artificially keeping bees and the disorientation of foragers. Despite the likely benefits of increased spacing between hives for bees, arranging nests many metres apart from one another may not be convenient or possible for beekeepers, especially with stingless bees, where the most common practice tends to be lining up the colonies just a few centimetres apart from one another (Leão et al., 2025; Nogueira-Neto, 1997; Pereira et al., 2017). Additionally, stingless beekeeping is often a small-scale practice conducted by communities and family-based groups (Carvalho et al., 2014; Costa et al., 2012; Jaffé et al., 2015; Leão et al., 2025; Martínez-Puc et al., 2024; Pereira et al., 2017), frequently with limited space and resources to keep hives in large infrastructures. Although maintaining colonies in proximity offers important logistical advantages, beekeepers frequently report problems that may be associated with drifting, including weakened colonies, particularly in the central area of the meliponary, marked variation in honey production, worker fighting, difficulties in queen replacement, and unusually large colony populations (Bruening, 1990; Koffler et al., 2015; Nogueira-Neto, 1997, CM, *personal communication*).

To better understand how hive arrangement affects drifting in stingless bees, we investigated the effect of different hive layouts on foraging activity and drifting behaviour in the stingless bee *Melipona quadrifasciata*, a common species in Brazilian meliponiculture (Jaffé et al., 2015) and an important pollinator of native plants (Barth et al., 2020; Wilms et al., 1996) and crops (Nogueira-Neto et al., 1959; Nunes-Silva et al., 2013; Viana et al., 2014). First, we analysed forager traffic in six layouts that varied in distance, colour and entrance orientation. Then, we used radio-frequency identification (RFID) tags to monitor forager drifting across three layouts with the same inter-hive distance, but different visual designs. Specifically, we investigated drifting behaviour in: i) visually similar hives, ii) hives with opposite entrance orientation, and iii) hives painted with distinct colours. To our knowledge, our study was the first to assess the effects of different hive layouts on drifting rates for a stingless bee species. We expected to find the highest drifting rates in the first layout, due to the lack of visual cues to differentiate the hives, and we also expected bees to drift more from central hives and to join their closest neighbour at higher rates.

## Material and Methods

### Study site and species

The study was conducted at the *Embrapa Meio Ambiente* meliponaries in Jaguariúna, São Paulo, Brazil. We performed experiments with the stingless bee *Melipona quadrifasciata,* a frequently managed (Jaffé et al., 2015; Nogueira-Neto, 1997) emerging crop pollinator in Brazil (Barth et al., 2020; Silva-Neto et al., 2019; Viana et al., 2014). Colonies of *M. quadrifasciata* have a typical size range of 300 to 1500 bees, with an average of 900 individuals (Grüter, 2020). In the first data collection period, between March and May 2025, we placed 30 colonies of *M. quadrifasciata* in wooden boxes (19 × 19 × 16 cm) in six different configurations (Figure 1).

**Figure 1.**
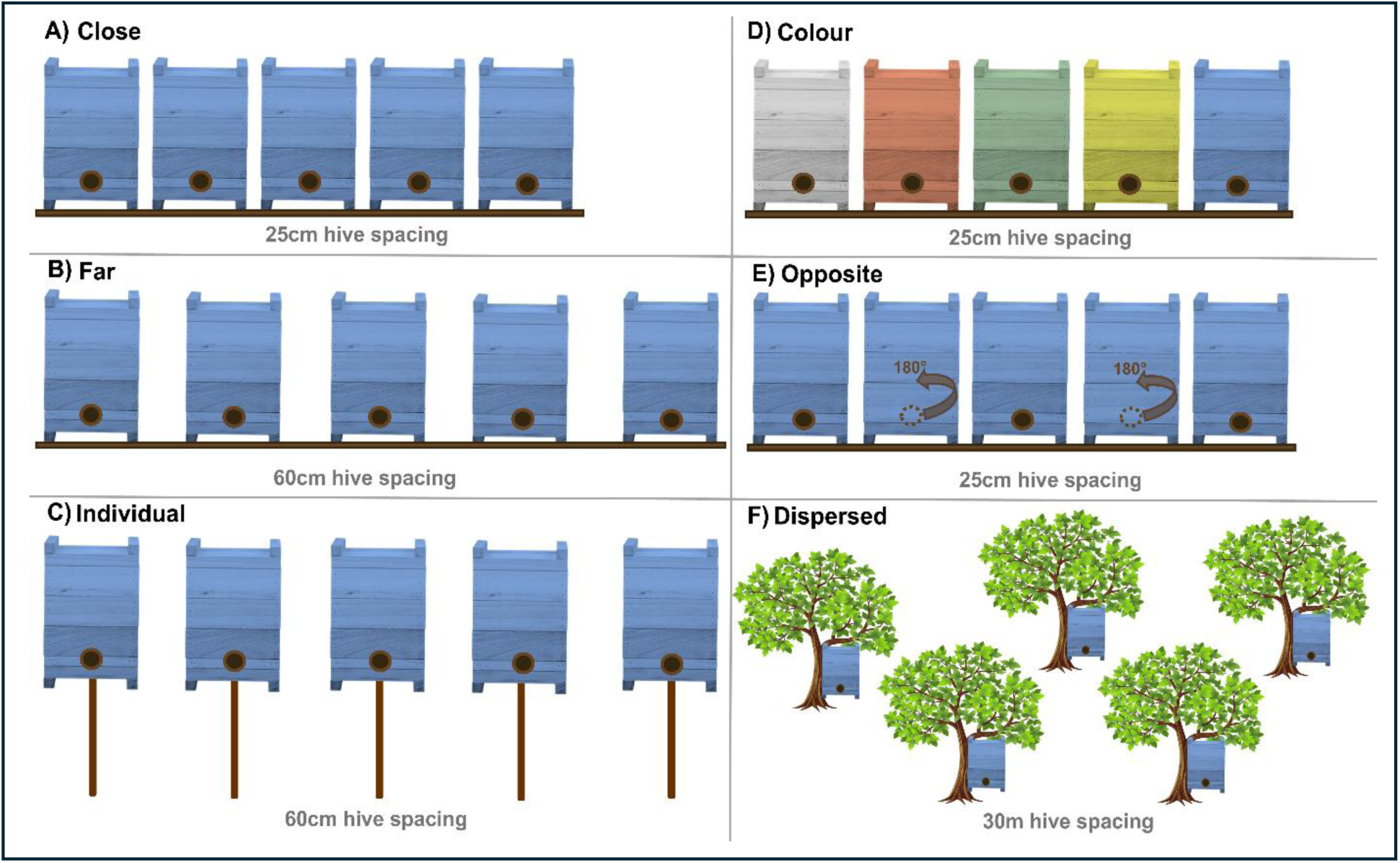
Hive configurations A) ‘Close’, in which blue hives were 25cm apart between the closest hive walls; B) ‘Far’, in which blue hives were 60cm apart; C) ‘Individual easel’, blue hives were placed 60cm apart on individual easels, a common practice with stingless bees; D) ‘Colour’, hives were painted with different colours (i.e. white, red, green, blue and yellow), and placed 25cm apart on a shelf; E) ‘Opposite’, blue hives were placed on a shelf 25cm apart with entrances facing opposite orientations, in a 180° (forward and backwards); F) ‘Dispersed’, blue hives were hung in dispersed trees in the environment, at least 30m apart from each other.

### Part 1: foraging activity in relation to hive layout

We allowed colonies to adapt for two weeks before we started monitoring forager activity. We had six different hive configurations (Figure 1), varying in the distance between hives, the arrangement, and the hive colour. The layout ‘close’ had blue hives 25cm apart (considering the closest hive walls) on a shelf; the layout ‘individual easel’ consisted of blue hives placed 60cm apart on individual easels, a common practice with stingless bees; the layout ‘colour’, in which hives were painted with different colours (i.e. white, red, green, blue and yellow), and placed 25cm apart on a shelf; the layout ‘opposite’ had blue hives placed on a shelf 25cm apart with entrances facing opposite orientations, in a 180° (forward and backwards); the layout ‘dispersed’ had blue hives hung in dispersed trees in the environment, at least 30m apart from each other.

To analyse the effects of apiary layout on foraging dynamics, we inspected forager traffic at the hive entrance (i.e., number of foragers), which is a non-invasive predictor of hive strength (Leão et al., 2025). To assess forager traffic, we registered the number of returning foragers for each hive for three minutes in three layouts per day, between 7 and 7:30 am in days with good weather conditions, for four consecutive weeks, and included a final monitoring in the ninth week.

### Part 2: drifting patterns in relation to hive layout

To study drifting patterns, data were collected between February and April 2026. Based on the results from Part 1, we selected three layouts to investigate associations between drifting rates and hive position. The selected arrangements were: ‘close’, ‘opposite’ and ‘colour’ (Figure 1), and each layout was arranged at the final location three months before the start of the drifting experiment. These three layouts were chosen due to the differences in flux during the first two weeks of the Part 1 experiment, in which bees from the ‘close’ layout had the lowest forager traffic, compared to the ‘colour’ ones (see Results section). Even though the layouts ‘far’ and ‘dispersed’ also had higher forager traffic compared to ‘close’, we wanted to test hive configurations that would be more convenient for beekeepers to inspect hives. Also, the configuration ‘close’ is the most used by professional beekeepers in Brazil (see Figure 2), whilst ‘colour’ and ‘opposite’ are also used, not as often as ‘close’, but these last two may improve foragers’ orientation and avoid aggression between hives (CM, *personal communication*).

**Figure 2.**
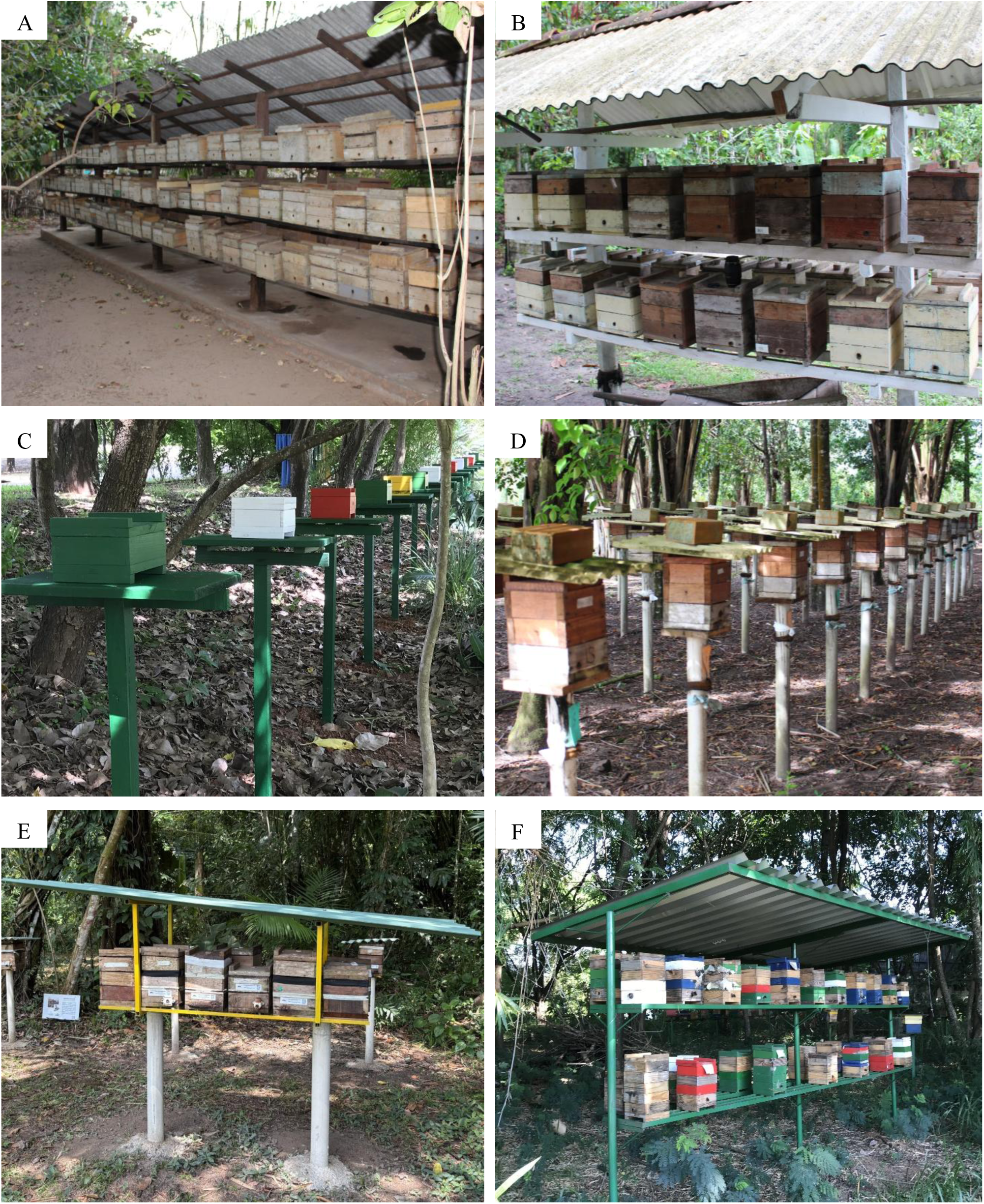
Photos (A-F) of meliponaries across Brazil, illustrating the diversity of hive layouts used by stingless beekeepers. (Photos: Cristiano Menezes).

Forager drifting was investigated using RFID tags (iID®BEEscience - MicroSensys, Magnay et al., 2025). Each bee was tagged with a mic3®Q1.6 RFID transponder (1.6 × 1.6 × 0.4 mm, 2.1 mg) and detected by iID®science readers (101 × 55 × 30 mm) fitted with dual antennae. The readers were attached at the hive entrance using screws and tape. This dual-antenna configuration allows the system to determine direction of travel (arriving or departing) with high directional accuracy (up to 93%, Magnay et al., 2025). Each layout was investigated for a period of two weeks, as a previous study with *Melipona fasciculata* bees using RFID technology recorded that their average activity after being tagged was 9.3 days (Oliveira et al., 2021). We randomly collected 80 foragers departing from each hive and used odourless cyanoacrylate adhesive to glue the RFID tags onto their thorax. Thus, we tagged 400 bees per layout and 1200 in total.

### Data Analysis

All statistical analyses were performed with R software (version 4.3.3). For the forager traffic data, we fitted zero-inflated negative binomial GLMMs, including layout and week, or hive position (i.e., edge, intermediate or centre) and layout, as fixed effects, with the interaction between these effects tested in each model. Colony identity was included as a random effect in all models. Fixed effects interactions were tested using Type III Wald χ² tests. For drifting information, the RFID data filtering and merging were performed using custom Python scripts (available here: <u>BeeDrifting</u>). The drifting probability across hive positions in different layouts was analysed using a binomial GLMM with the interaction between hive position and layout as the fixed effect and hive identity as a random effect. Also, we used binomial GLMMs to investigate drifting direction, considering hives within the same layout as targets and categorising each drifting event as visited (1) or not (0). The fixed effects were ‘distance class’: 1^st^ neighbour (1-hive distance), 2^nd^ neighbour (2-hive distance) and distant neighbour (3 or 4-hive distance), and its interaction with layout. Hive and bee IDs were included as random effects. For layout and distance-class contrasts based on a few visits, we additionally applied Firth’s bias-reduced logistic regression as a robustness check on the estimates of the fixed effects. Across all models, we tested the significance of fixed effects and their interaction with Type III Wald χ² tests.

We also evaluated the number of foreign hives visited by tagged foragers in each layout and classified them into three behavioural categories: ‘resident foragers’ were bees that were never detected at a hive other than their hive of origin. Among drifters, those recorded at a single foreign hive and never subsequently detected back at their original hive were classified as ‘migrated foragers’, indicating a one-way relocation. Foragers that either returned to their original hive after visiting a foreign hive or were detected at more than one foreign hive were classified as ‘transient foragers’, reflecting more extensive movement between colonies.

## Results

### Part 1: forager traffic in relation to hive layout

We found interactions between layout and week (GLMM, *χ²* = 93.58, *df* = 20, *p* < 0.001). In the first week, bees in the ‘close’ layout had lower forager traffic in relation to the ones in the ‘colour’ layout (GLMM, *z* = −3.82, *p* = 0.002, Figure 3), representing a 6.4-fold reduction for hives positioned close (i.e., visually similar and 25cm apart). We also found a 4.3-fold lower foraging activity when comparing the ‘close’ layout to ‘far’ (GLMM, *z* = −2.91, *p* = 0.025) and to the ‘dispersed’ layout (GLMM, *z* = −2.804, *p* = 0.025), with a 3.9-fold reduction. We found no differences between the ‘close’ and ‘individual’ layouts (GLMM, *z* = −2.35, *p* = 0.071) or opposite (GLMM, *z* = −1.50, *p* = 0.288). In the second week, foragers from the ‘close’ layout also had a lower foraging activity compared to the ones in the ‘colour’ layout (GLMM, *z* = −3.05, *p* = 0.034, Figure 3), representing a 4.5-fold reduction. Bees in the ‘individual’ layout had 3.7 times lower external activity in relation to the ‘colour’ layout (GLMM, *z* = −2.74, *p* = 0.046). On the other hand, we found no differences between layouts in the third, fourth or ninth weeks of experiments (GLMM, *|z|* < 2.46, *p* > 0.21).

**Figure 3.**
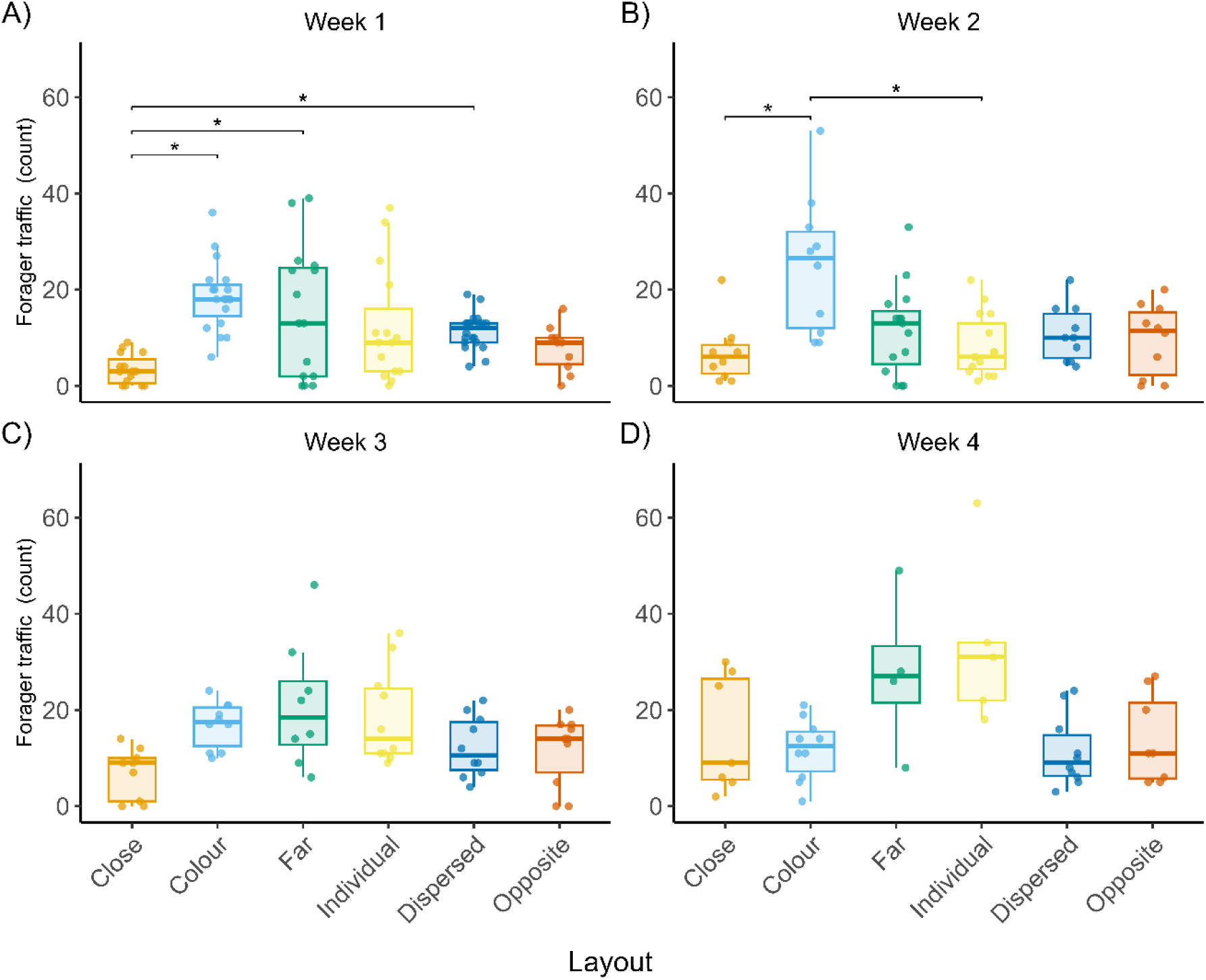
Forager traffic per layout across weeks. A) Week 1, B) Week 2, C) Week 3, D) Week 4. Boxes represent the median and interquartile range (IQR); individual points represent forager count per hive. Asterisks (*) indicate significant differences between layouts (p < 0.05).

We found an interaction between hive position and layout (GLMM, *χ²* = 25.03, *df* = 10, *p* = 0.005) on forager activity. In the ‘far’ layout, bees in the central hive had the lowest external activity compared to intermediate (GLMM, *z* = −4.44, *p* < 0.0001) and edge hives (GLMM, z = −3.68, *p* = 0.0004). However, in the ‘close’, ‘opposite’, ‘dispersed’, ‘individual’ and ‘colour’ layouts, we found no differences (GLMM, *|z|* < 1.60, *p* > 0.23).

### Part 2: forager drifting in relation to hive layout

Drifting rates ranged from 37.6% in the ‘colour’ layout, to 48.1% in the ‘opposite’ layout and 59.5% in the ‘close’ layout, indicating that over one-third to nearly two-thirds of tagged bees drifted at least once. Specifically, over the two-week monitoring period per layout, bees from edge hives drifted less, in a range of 16.4% to 18.3%, compared to 52.8% to 91% from the centre hives, and 51.1% to 86.5% from the intermediate hives.

We found an interaction between layout and hive position on the drifting proportion (*χ²* = 16.78, *df* = 2, *p* < 0.001). In the ‘close’ layout, the drifting probability from the inner hives was 88.0%, compared to 16.4% from the edge hives (GLMM, *z* = 10.06, *p* < 0.0001; Figure 4). In the ‘opposite’ layout, the drifting probability was 68.8% from the inner hives and 18.3% from the edge hives (GLMM, *z* = 6.98, *p* < 0.0001). In the ‘colour’ layout, the drifting probability was also higher in the inner positions, with 51.7% from inner hives and 16.6% from the edge (GLMM, *z* = 5.11, *p* < 0.0001). Thus, the proportion of drifting for each layout depended on hive position, with edge hive bees drifting considerably less than those from inner positions across all layouts (Figure 4). Considering the inner hives, drifting was 3.4 times higher in the ‘close’ layout compared to the ‘opposite’ layout (GLMM, *z* = 3.93, *p* = 0.0002), 7.0 times higher compared to the ‘colour’ layout (GLMM, *z* = 6.38, *p* < 0.0001), and 2.1 times higher in the ‘opposite’ layout compared to the ‘colour’ layout (GLMM, *z* = 2.69, *p* = 0.0072). At edge hives, drifting rates did not differ between layouts (GLMM, *|z|* < 0.32, *p* = 1.000, Figure 4).

**Figure 4.**
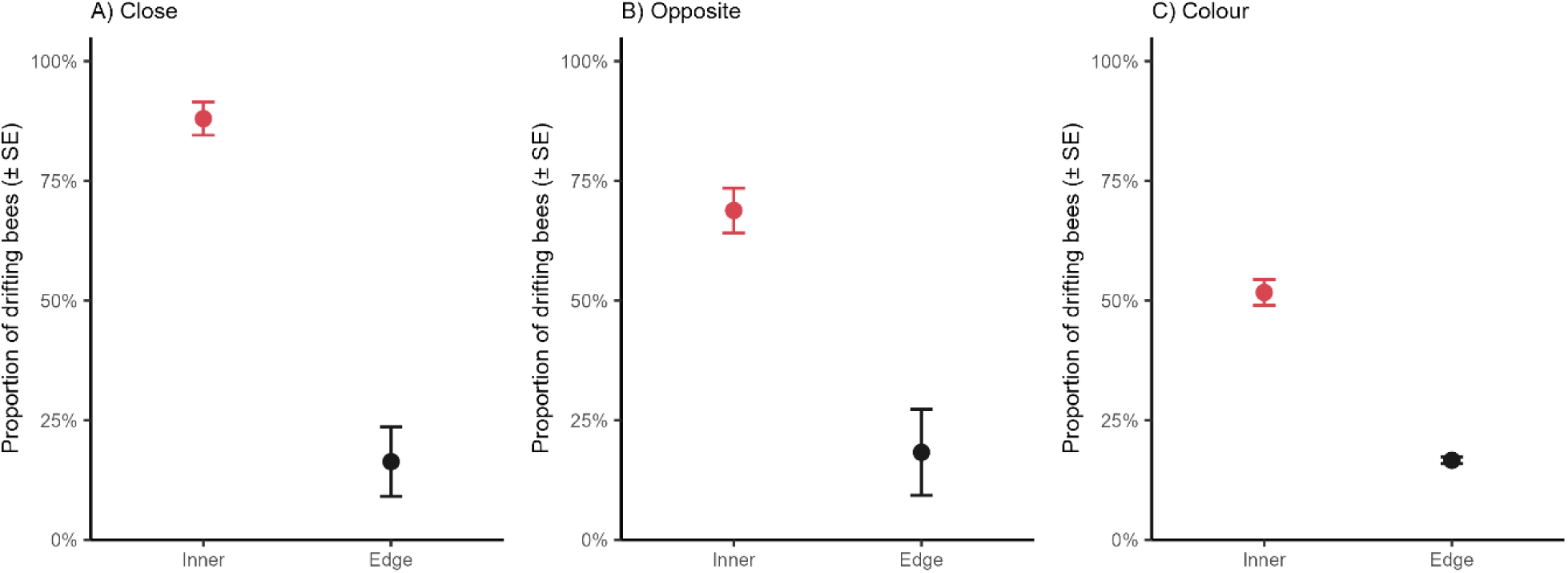
Mean (± SE) proportion of drifting bees according to hive position within each layout: A) Close, B) Opposite, C) Colour: Red dots represent inner hives (centre + intermediates) and black dots represent edge hives.

We found that foragers drift to neighbouring hives at different rates, according to the layout and distance class (GLMM, *χ²* = 162.92, *df* = 4, *p* < 0.001; Figure 5). In the ‘close’ layout, bees had a higher probability of drifting to neighbouring hives (i.e., 1^st^ and 2^nd^ neighbours) than to the most distant ones (GLMM, 1^st^ vs distant: *z* = 8.65, *p* < 0.0001; 2^nd^ vs distant: *z* = 9.33, *p* < 0.0001; 2nd vs 1st: *z* = −1.97, *p* = 0.049). In the ‘opposite’ layout, the majority of visits were directed to the 2^nd^ neighbour hives, compared to the 1^st^ neighbours (GLMM, *z* = 9.69, *p* < 0.0001) or to the distant ones (GLMM, 2^nd^ vs distant: *z* = 6.17, *p* < 0.0001), while 1^st^ and distant neighbours did not differ (GLMM, *z* = 0.35, *p* = 0.727). In the ‘colour’ layout, the majority of drifting was to the 1^st^ neighbour hives compared to the 2^nd^ (GLMM, *z* = 6.79, *p* < 0.0001) and distant (GLMM, *z* = 6.02, *p* < 0.0001), while 2^nd^ and distant neighbours did not differ (GLMM, *z* = 0.66, *p* = 0.505). Lastly, Firth’s regression confirmed the same patterns for the contrasts investigated, except for the opposite layout 1^st^ vs distant contrast, as the visits were too few (1^st^ neighbour: 3 visits, distant neighbour: 1 visit).

**Figure 5.**
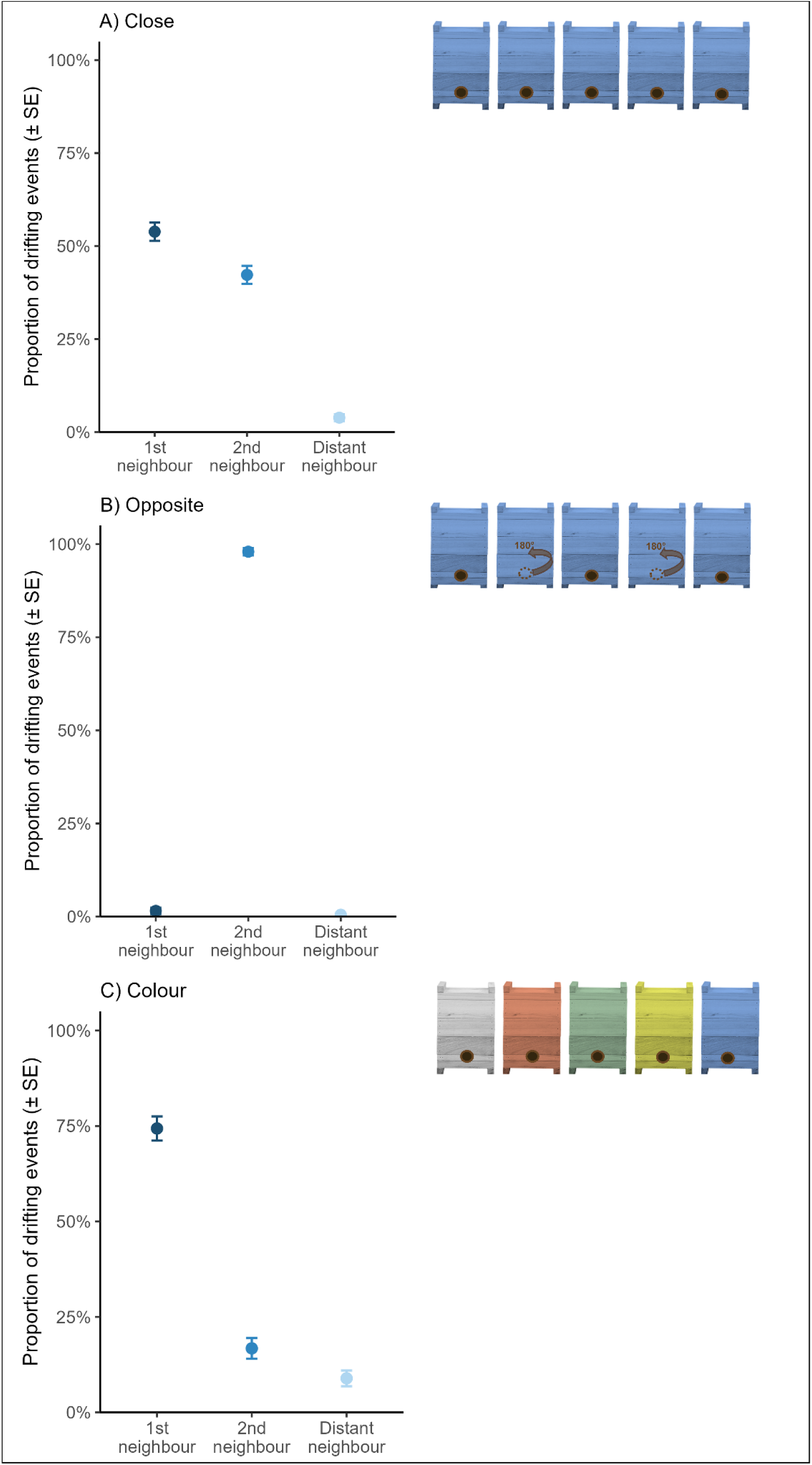
Mean (± SE) proportion of drifting events by distance class from the natal hive (1st neighbour, 2nd neighbour, distant neighbour), for each layout: A) Close, B) Opposite, C) Colour, with schematics of each arrangement shown above. Point shading reflects this same distance gradient, from dark blue (1st neighbour) to light blue (distant neighbour).

Additionally, we found that drifting dynamics varied according to the layout and hive position, both in the number of foreign hives visited, with bees from ‘close’ hives drifting to a higher number of foreign hives and at a greater proportion (Figure 6A), and in the proportion of bees classified as ‘resident’, ‘migrated’, or ‘transient’ (Figure 6B).

**Figure 6.**
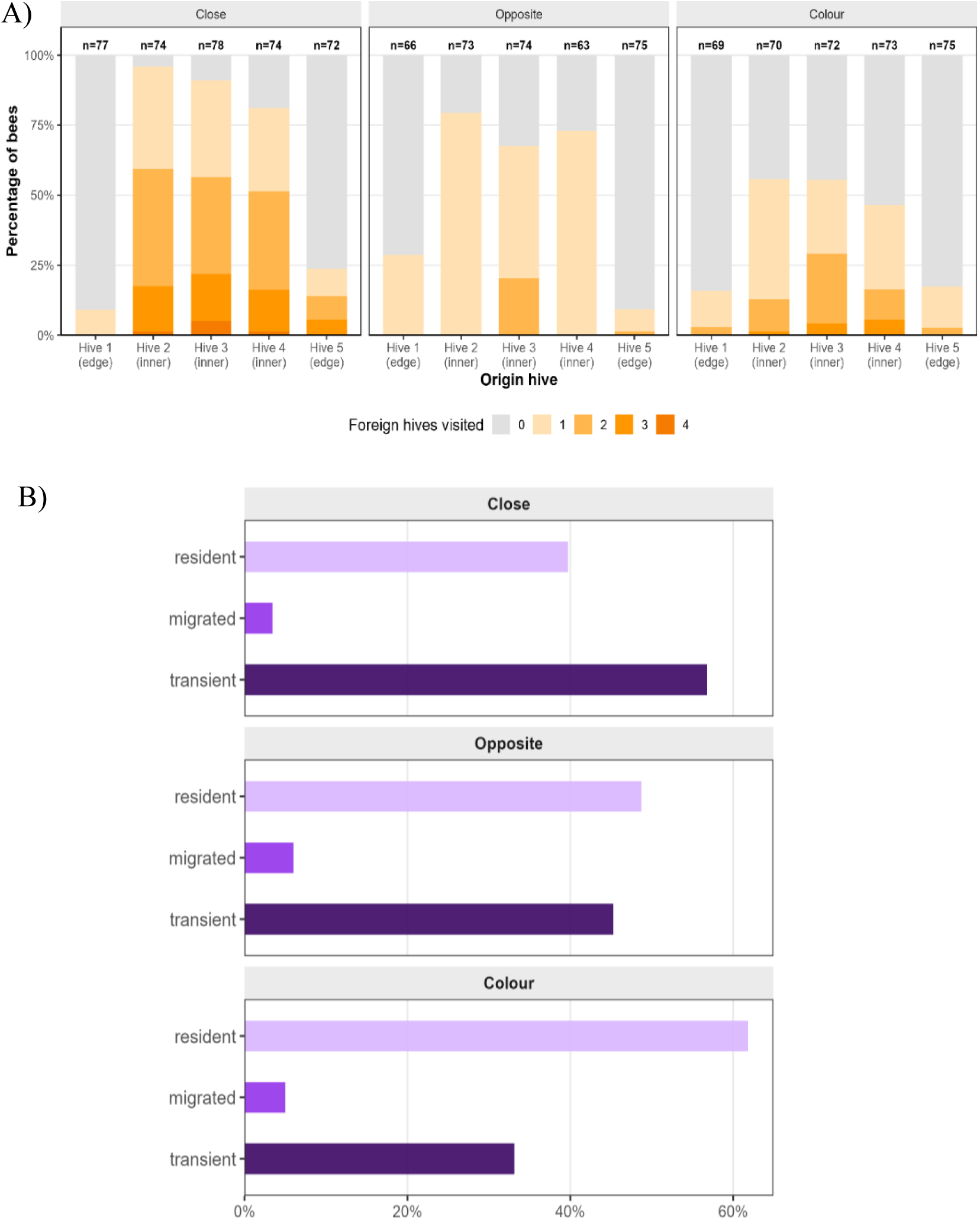
Drifting dynamics. A) Percentage of bees per origin hive according to the number of foreign hives visited. Colours represent the number of foreign hives visited (0 to 4). N indicates the total number of bees recorded per hive. B) Proportion of bees classified as resident, migrated, or transient across the study population.

## Discussion

Forager traffic was affected by hive layout during the first two weeks of monitoring. Bees from the ‘close’ layout had the lowest traffic compared to bees in layouts where hives had different colours or were placed at greater distances. Interestingly, drifting rates were higher in the ‘close’ layout compared to the ‘colour’ and ‘opposite’ layouts, with bees from inner hives driving most of these differences. Hive position (inner or edge) had a substantial effect on drifting rates, while layout affected the proportion of bees that drifted, the number of foreign hives visited, and the most commonly visited neighbour hive.

The differences in forager traffic are likely associated with bee disorientation during the initial adaptation period. This disorientation is expected to be greater in the ‘close’ layout, in which bees lacked clear cues that could have increased their ability to recognise their original hive. In contrast, in the ‘colour’, ‘far’, and ‘dispersed’ layouts, visual cues (i.e., hive colour and distance) likely improved the rate of bees returning to the right colony, resulting in fewer foragers getting lost. This probably reduced aggressive conflicts with non-nestmates, which have been reported in *M. quadrifasciata* bees (Nogueira-Neto, 1997). When leaving a nest for the first time, bees usually turn and hover for a few seconds in front of the nest entrance, performing repeated looping flights of progressively increasing radius before departing in a straight line (Roubik, 1989; Zeil et al., 1996). In honeybees (*Apis mellifera*), these orientation flights are known to play a crucial role in learning the location of the hive and its surroundings (Capaldi and Dyer, 1999; Degen et al., 2016; Wei and Dyer, 2009). In this process, foragers observe the environment and learn the landscape characteristics to guide themselves during foraging trips (Capaldi and Dyer, 1999; Degen et al., 2016), and evidence suggests that even a single orientation flight provides enough landscape information about the vicinity of the hives to guide bee navigation (Capaldi et al., 1999; Degen et al., 2016). In this sense, high disorientation and drifting are expected in high-density apiaries with visually similar hives. Sharing a neighbourhood with visually similar hives is not a natural spatial configuration for bees (Seeley, 2007; Stephens et al., 2017), which likely lack biological mechanisms to discriminate hives positioned a few centimetres apart and similar in appearance.

Forager disorientation and drifting have previously been shown to increase when honeybee hives were newly introduced to a location (Nelson and Jay, 1989). Additionally, RFID data from *Melipona seminigra* bees show a higher proportion of disoriented foragers following relocation, with homing behaviour normalising after one or two months in the new location (Costa et al., 2021). These patterns of disorientation and drifting could explain the differences in forager traffic we found here. Bees in the visually similar layout (i.e., ‘close’) may have experienced greater disorientation during the first two adaptation weeks, resulting in greater loss through aggressive encounters with non-nestmate bees from neighbouring hives. We observed fights between guards and forager bees during this adaptation period (APC, *personal observation*). Such aggressive encounters were not observed during the second experiment, in which hives from each layout shared the same location for at least three months prior to the start of the observations.

Stingless bees are reported to engage in aggressive intraspecific conflicts that can result in high rates of forager mortality (Bruening, 1990; Cunningham et al., 2014; Grüter, 2020; Hubbell et al., 1977; Johnson and Hubbell, 1974; Nogueira-Neto, 1997). For example, in some *Trigona* species, intraspecific aggression plays a role when bees are attempting to build their nests close to pre-established hives (Hubbell and Johnson, 1977). Hubbell and Johnson (1977) suggest that natural spacing between hives can be actively maintained through aggressive exclusion, associating hive proximity with costly conflicts. In meliponaries, some bee species are known to not tolerate the presence of others (Nogueira-Neto, 1997). In *Melipona subnitida,* aggression between hives positioned on shelves has been reported, with large numbers of bees involved in fatal fights lasting several days (Bruening, 1990). Nogueira-Neto (1997), specifically for *M. quadrifasciata*, reported robbing and fights between neighbouring hives within meliponaries, describing that this aggressiveness problem was largely resolved by increasing the space between hives and changing their orientation. Additionally, differences in guard rejection rates can also influence foreign bees visits and aggression rates in meliponaries; for example, *Tetragonisca angustula* guards are extremely precise in recognising non-nestmates (i.e., a 92% rejection rate) (Kärcher and Ratnieks, 2009), and often engage in fatal fights. In *M. quadrifasciata*, guards discriminated against and attacked 74% of non-nestmate conspecifics presented to them, contrasting with the rejection rates of *M. scutellaris* and *M. rufiventris* (14% and 60%, respectively) (Breed and Page, 1991). More recently, Batista and collaborators (2024) showed that *M. quadrifasciata* hives kept in proximity rejected nestmates and non-nestmates at similar rates, potentially indicating chemical homogenisation via drifting and a reduced ability of guards to discriminate drifters from non-drifters. Thus, hive layouts that favour forager disorientation can impact forager loss, disease spread, and hive productivity.

In our study, the opposite entrance orientation and the distinct colour layouts reduced the overall drifting proportion compared to the visually similar layout (‘close’). However, we found that bees almost never (less than 1%) drifted to hives with an entrance positioned in the opposite direction of their original hive. This information can be used to improve colony health and mitigate the impacts of drifting in managed hives, particularly because each of our layouts included five hives and already showed high drifting rates. These effects are likely to be amplified in commercial meliponaries with up to 100 hives placed in close proximity. While studies with honeybees demonstrated a decrease in drifting when hives were arranged in distinct configurations (Dynes et al., 2019; Seeley and Smith, 2015) these layouts tested a great contrast in distance (e.g., 1m vs 10m or >30m), and differences in hive colour, height and position in the space (Dynes et al., 2019), which makes it difficult to disentangle the effects of the different factors (i.e., colour, orientation and distance). In the stingless bee *M. fasciculata,* Oliveira and collaborators (2021) found an increase in drifting rates when geometrical patterns were positioned at hive entrances, which contrasts with our findings and might indicate other factors affecting drifting rates in *Melipona* bees, such as robbing or parasitism (Blacher et al., 2013; Grüter, 2020; Peck and Seeley, 2019). However, in our results, the majority of bees visited the 1^st^ neighbour in the ‘close’ and ‘colour’ layouts, and the 2^nd^ neighbour in the ‘opposite’ layout, which suggests bees are joining foreign hives that are spatially close and in the same orientation as their original hive, indicating forager disorientation in an artificial hive context. These results corroborate the drifting patterns found in a modelling environment where bee drifting was an emergent property of navigational mistakes, with greater drifting rates at small distances and bees drifting more from inner hives compared to edge, which influenced hive population and energy (Cipriano et al., 2026). Overall, our results with *M. quadrifasciata* show that hive layouts are associated with considerable differences in foraging and drifting behaviour, an important aspect to be considered for colony health and honey production.

*Melipona* hives often show considerable discrepancies in inter-hive honey production, even when the hives are allocated in standardised managed boxes and in the same type of environment (Farfan et al., 2023; Kerr et al., 2001; Koffler et al., 2015; Ostrovski, 2019). For example, in the same environmental conditions and type of boxes, *M. subnitida* honey production varied from 0g to 1,800g in the same year (Koffler et al., 2015), and a similar discrepancy has also been described for *M. quadrifasciata* bees (Ostrovski, 2019). Substantial inter-hive productivity discrepancies should be avoided in beekeeping, as unproductive hives may increase the management effort, e.g. by artificial feeding or the addition of brood to strengthen worker populations. In managed hives, differences in drifting rates across hive layouts contribute to variation in honey production (Dynes et al., 2019). In *A. mellifera*, when comparing high-density and low-density apiaries, Dynes and collaborators (2019) found differences in drifting rates and honey production, and the low-density apiary (i.e., hives where 10m apart and with visual cues) produced significantly more honey, with bees drifting 3 times less. In addition, crowded apiaries also contributed to amplifying parasite infection between hives and reduced the rate of winter survival in infected hives (Dynes et al., 2019). Thus, hive layouts that reduce drifting are likely to affect not only honey productivity, but also health and survival in social bees.

The impact of pests and diseases on honey production and colony loss is an alarming issue for both honeybee and stingless beekeepers (Dias de Freitas et al., 2023). Particularly for stingless bees, the complementary feeding with honeybee pollen or honey increases the parasite load within hives (Dias de Freitas et al., 2023; Grüter, 2020), which may contribute to hive contamination. Additionally, pathogen spillover on flowers visited by *A. mellifera*, for instance, has been shown to contribute to *Nosema ceranae* contamination in *Tetragonula hockingsi* (Purkiss and Lach, 2019). This pathogen can reduce bee longevity and impact foraging (Purkiss and Lach, 2019), with older foragers drifting more and performing shorter trips when *A. mellifera* were infected with *N. ceranae* (Bordier et al., 2017), which likely contributes to the pathogen spread between neighbouring hives. Other diseases that are extremely detrimental to hives, causing brood loss and colony deaths, such as European foulbrood, have recently been detected in Brazilian stingless bees, with several species infected in distinct locations, including *M. quadrifasciata* bees (Teixeira et al., 2025). Drifting bees are vectors of pathogens (Retschnig et al., 2019; Šekulja et al., 2014), in *A. mellifera,* for example, Varroa-infested hives accepted drifters at a higher rate (Forfert et al., 2015), and bees infected with Israeli acute paralysis virus were more accepted by foreign guards (Geffre et al., 2020), which potentially increases parasite and pathogen spread within the apiary. This combination of multiple environmental and spatial factors that risk pathogen spread and reduce colony productivity could act as important causes of stingless bee health problems. Exploring alternatives to reduce drifting is therefore a promising area for mitigating these concerns.

The increasing importance of stingless bees for crop pollination (Nunes-Silva et al., 2013; Silva-Neto et al., 2019; Silva et al., 2015; Slaa et al., 2006), resource commercialisation (Imperatriz-Fonseca et al., 2017; Jaffé et al., 2015), and for cultural and environmental reasons (Carvalho et al., 2014; Grüter, 2020; Martínez-Puc et al., 2024; Quezada-Euán, 2018) highlights the importance of understanding environmental factors that impact their health and survival. Our results highlight the importance of investigating how to improve meliponary layout and, consequently, reduce the risks of forager loss, pathogen contamination, population imbalance, and reduced honey production. Our findings suggest that alternating hive entrances and including visual cues such as different colours substantially reduce drifting rates and potentially improve hive health and productivity. Future research should aim to further understand meliponary arrangements that reduce drifting without increasing the management effort, for example, by alternating hives of different species that tolerate each other. Also, research should integrate the impact of stressors such as pesticides, pathogens, and low-quality food sources on bee drifting. Thus, research investigating the behaviour and ecology of stingless bees, a group that is understudied compared to honeybees and bumblebees, also helps shed light on the biology of these native pollinators, expanding our understanding of their foraging patterns.

## Acknowledgements

We would like to thank Embrapa Meio Ambiente staff for their technical support and Mariana F. Santos for her help in forager traffic data collection. A.P.C. was funded by South West Biosciences Doctoral Training Partnership (SWBio DTP) (BB/T008741/1), and M.M. was funded by the University of Bristol PGR fund.

## Bibliography

Barth, O. M., Freitas, S. De, Vanderborght, B., Monika, O., Freitas, S. De, & Vanderborght, B. (2020). Pollen preference of stingless bees (Melipona rufiventris and M. quadrifasciata anthidioides) inside an urban tropical forest at Rio de Janeiro city. Journal of Apicultural Research, 59(5), 1005–1010. doi: 10.1080/00218839.2020.1714863

Batista, J. E., da Silva, R. C., do Nascimento, D. L., Oliveira, R. C., Oi, C. A., & do Nascimento, F. S. (2024). Nestmate Recognition in Two Melipona Stingless Bee Species: The Effect of Cuticular Chemical Profiles and Colony Distance. Journal of Insect Behavior, 37(1), 106–120. doi: 10.1007/s10905-024-09852-z

Blacher, P., Yagound, B., Lecoutey, E., Devienne, P., Chameron, S., & Châline, N. (2013). Drifting behaviour as an alternative reproductive strategy for social insect workers. Proceedings of the Royal Society B: Biological Sciences, 280(1771). doi: 10.1098/rspb.2013.1888

Bordier, C., Pioz, M., Crauser, D., Le Conte, Y., & Alaux, C. (2017). Should I stay or should I go: honeybee drifting behaviour as a function of parasitism. Apidologie, 48(3), 286–297. doi: 10.1007/s13592-016-0475-1

Breed, M., & Page, R. (1991). Intra- and interspecific nestmate recognition in Melipona workers (Hymenoptera: Apidae). Journal of Insect Behavior, 4(4), 463–469. doi: 10.1007/BF01049331

Bruening, M. H. (1990). Abelha jandaira. Coleção Mossoroense (RN), série C, v.557, (DIYI1).

Bueno, F. G. B., Kendall, L., Alves, D. A., Tamara, M. L., Heard, T., Latty, T., & Gloag, R. (2023). Stingless bee floral visitation in the global tropics and subtropics. Global Ecology and Conservation, 43, e02454. doi: 10.1016/j.gecco.2023.e02454

Capaldi, E. A., & Dyer, F. C. (1999). The role of orientation flights on homing performance in honeybees. Journal of Experimental Biology, 202(12), 1655–1666. doi: 10.1242/jeb.202.12.1655

Carvalho, R. M. A. de, & Martins, C. F. (2014). É Uma Abelha Sagrada”: Dimensão Simbólica Da Criação De Abelhas Sem Ferrão Em Comunidades Quilombolas Da Zona Da Mata Sul Paraibana. Gaia Scientia, 15–27. Retrieved from http://periodicos.ufpb.br/ojs/index.php/gaia/article/view/22405

Cipriano, A. P., Bell, E., & Grüter, C. (2026). Drifting behaviour in social bees: using agent-based modelling to explore the effects of spatial layout on drifting patterns. Available at SSRN 6983299.

Cortopassi-Laurino, M., Imperatriz-Fonseca, V. L., Roubik, D. W., Dollin, A., Heard, T., Aguilar, I., Venturieri, G. C., Eardley, C., & Nogueira-Neto, P. (2006). Global meliponiculture: challenges and opportunities. Apidologie, 37, 275–292.

Costa, L., Nunes-Silva, P., Galaschi-Teixeira, J. S., Arruda, H., Veiga, J. C., Pessin, G., de Souza, P., & Imperatriz-Fonseca, V. L. (2021). RFID-tagged amazonian stingless bees confirm that landscape configuration and nest re-establishment time affect homing ability. Insectes Sociaux, 68(1), 101–108. doi: 10.1007/s00040-020-00802-4

Costa, T. V., Farias, C. A. G., & Brandão, C. dos S. (2012). Meliponicultura em comunidades tradicionais do Amazonas. Revista Brasileira de Agroecologia, 7(3), 106–115.

Cunningham, J., Hereward, J., Heard, T., De Barro, P., & West, S. (2014). Bees at war: interspecific battles and nest usurpation in stingless bees. Am Nat, 184, 777–786.

Degen, J., Kirbach, A., Reiter, L., Lehmann, K., Norton, P., Storms, M., Koblofsky, M., Winter, S., Georgieva, P. B., Nguyen, H., Chamkhi, H., Meyer, H., Singh, P. K., Manz, G., Greggers, U., & Menzel, R. (2016). Honeybees Learn Landscape Features during Exploratory Orientation Flights. Current Biology, 26(20), 2800–2804. doi: 10.1016/j.cub.2016.08.013

Dias de Freitas, C., Oki, Y., Resende, F. M., Zamudio, F., Simone de Freitas, G., Moreira de Rezende, K., Amaro de Souza, F., De Jong, D., Quesada, M., Siqueira Carvalho, A., Silvia Soares Pires, C., & Fernandes, G. W. (2023). Impacts of pests and diseases on the decline of managed bees in Brazil: a beekeeper perspective. Journal of Apicultural Research, 62(5), 969–982. doi: 10.1080/00218839.2022.2099188

Dynes, T. L., Berry, J. A., Delaplane, K. S., Brosi, B. J., & de Roode, J. C. (2019). Reduced density and visually complex apiaries reduce parasite load and promote honey production and overwintering survival in honey bees. PLoS ONE, 14(5), e0216286. doi: 10.1371/journal.pone.0216286

Engel, M. S., Rasumussen, C., Ayala, R., & de Oliveira, F. F. (2023). Stingless bee classification and biology (Hymenoptera, Apidae): a review, with an updated key to genera and subgenera. ZooKeys, 1172, 239–312. doi: 10.3897/┘zookeys.1172.104944

Farfan, S. J. A., Celentano, D., Silva Junior, C. H. L., de Freitas Silveira, M. V., Serra, R. T. A., Gutierrez, J. A. M., Barros, H. C., Ribeiro, M. H. M., Barth, O. M., de Oliveira Alves, R. M., García, L. M. H., & Rousseau, G. X. (2023). The effect of landscape composition on stingless bee (Melipona fasciculata) honey productivity in a wetland ecosystem of Eastern Amazon, Brazil. Journal of Apicultural Research, 62(5), 1102–1114. doi: 10.1080/00218839.2022.2137307

Forfert, N., Natsopoulou, M. E., Frey, E., Rosenkranz, P., Paxton, R. J., & Moritz, R. F. A. (2015). Parasites and pathogens of the honeybee (Apis mellifera) and their influence on inter-colonial transmission. PLoS ONE, 10(10), 1–14. doi: 10.1371/journal.pone.0140337

Free, J. B. (1958). The drifting of honey-bees. The Journal of Agricultural Science, 51(3), 294–306. doi: 10.1017/S0021859600035103

Geffre, A. C., Gernat, T., Harwood, G. P., Jones, B. M., Gysi, D. M., Hamilton, A. R., Bonning, B. C., Toth, A. L., Robinson, G. E., & Dolezal, A. G. (2020). Honey bee virus causes context-dependent changes in host social behavior. Proceedings of the National Academy of Sciences of the United States of America, 117(19), 10406–10413. doi: 10.1073/pnas.2002268117

Giannini, T. C., Boff, S., Cordeiro, G. D., Cartolano, E. A., Veiga, A. K., Imperatriz-Fonseca, V. L., & Saraiva, A. M. (2015). Crop pollinators in Brazil: a review of reported interactions. Apidologie, 46(2), 209–223. doi: 10.1007/s13592-014-0316-z

Grüter, C. (2020). Stingless bees: their behaviour, ecology and evolution. Springer Nature.

Hubbell, S., & Johnson, L. (1977). Competition and Nest Spacing in a Tropical Stingless Bee Community. Ecology, 58(5), 949–963. Retrieved from https://www.jstor.org/stable/1936917

Imperatriz-fonseca, V. L., Koedam, D., & Hrncir, M. (2017). A abelha jandaíra no passado, no presente e no futuro. EdUFERSA.

Jaffé, R., Pope, N., Carvalho, A. T., Maia, U. M., Blochtein, B., Alfredo, C., Carvalho, L. De, & Carvalho-zilse, G. A. (2015). Bees for Development: Brazilian Survey Reveals How to Optimize Stingless Beekeeping. PLoS ONE, 1–21. doi: 10.1371/journal.pone.0121157

Jay, S. C. (1966). Drifting of Honeybees in Commercial Apiaries. III. Effect of Apiary Layout. Journal of Apicultural Research, 5(3), 137–148. doi: 10.1080/00218839.1966.11100147

Johnson, L., & Hubbell, S. (1974). Aggression and Competition among Stingless Bees: Field Studies. Ecology, 55(1), 120–127. Retrieved from https://www.jstor.org/stable/1934624

Kärcher, M., & Ratnieks, F. L. W. (2009). Standing and hovering guards of the stingless bee Tetragonisca angustula complement each other in entrance guarding and intruder recognition. Journal of Apicultural Research, 48(3), 209–214. doi: 10.3896/IBRA.1.48.3.10

Kerr, W. E., Petrere Jr, M., & Diniz Filho, J. A. F. (2001). Informações biológicas e estimativa do tamanho ideal da colmeia para a abelha tiúba do Maranhão (Melipona compressipes fasciculata Smith-Hymenoptera, Apidae). Revista Brasileira de Zoologia, 18(1), 45–52.

Koffler, S., Menezes, C., Menezes, P. R., Kleinert, A. D. M. P., Imperatriz-Fonseca, V. L., Pope, N., & Jaffé, R. (2015). Temporal Variation in Honey Production by the Stingless Bee Melipona subnitida (Hymenoptera: Apidae): Long-Term Management Reveals Its Potential as a Commercial Species in Northeastern Brazil. Journal of Economic Entomology, 108(3), 858–867. doi: 10.1093/jee/tov055

Leão, K. L., Campbell, A. J., Veiga, J. C., Andrés, F., Contrera, L., Leão, K., Campbell, A. J., & Veiga, J. C. (2025). Colony size of amazonian stingless bees and its assessment through intrinsic parameters intrinsic parameters. Journal of Apicultural Research, 64(3), 981–990. doi: 10.1080/00218839.2024.2327114

Leão, K., Menezes, C., Motta, M., & Aleixo, K. (2025). Curso de Meliponicultura para Mulheres. FAO. doi: ISBN 978-92-5-139842-5

Lepeco, A., Branstetter, M. G., Melo, G. A. R., Freitas, F. V., Tobin, K. B., Gan, J., Jensen, J., & Almeida, E. A. B. (2024). Phylogenomic insights into the worldwide evolutionary relationships of the stingless bees (Apidae, Meliponini). Molecular Phylogenetics and Evolution, 201, 108219. doi: 10.1016/j.ympev.2024.108219

Magnay, A. J., Cipriano, A. P., Versteeg, S. M., & Segers, F. (2025). Comprehensive Analysis of RFID Performance of the iID ® BEEscience system. BioRxiv, 1–24.

Martínez-Puc, J. F., Magaña-Magaña, M. Á., Cetzal-Ix, W., Mendoza-Arroyo, G. E., Sierra-Vasquez, Á. C., & Basu, S. K. (2024). Sociodemographic characteristics and participation of women in meliponiculture from the Yucatán Peninsula, Mexico. Journal of Ethnobiology and Ethnomedicine, 20(1), 1–11. doi: 10.1186/s13002-024-00745-1

Nelson, D., & Jay, S. (1989). Original article The effect of colony relocation of honeybees. Api, 20, 245–250.

Nogueira-Neto, P. (1997). Vida e criação de abelhas indígenas sem ferrão. São Paulo: Nogueira-Neto.

Nogueira-Neto, P., Carvalho, A., & Antunes, H. F. (1959). Efeito da exclusão dos insetos polinizadores na produção do café Bourbon. Bragantia, 18(29).

Nunes-Silva, P., Hrncir, M., Silva, I. C., & Imperatriz-Fonseca, V. L. (2013). Stingless bees, Melipona fasciculata, as efficient pollinators of eggplant (Solanum melongena) in greenhouses. Apidologie, 537–546. doi: 10.1007/s13592-013-0204-y

Oliveira, R. C., Contrera, F. A. L., Arruda, H., Jaffé, R., Costa, L., Pessin, G., Venturieri, G. C., de Souza, P., & Imperatriz-Fonseca, V. L. (2021). Foraging and drifting patterns of the highly eusocial neotropical stingless bee Melipona fasciculata assessed by radio-frequency identification tags. Frontiers in Ecology and Evolution, 9, 1–10. doi: 10.3389/fevo.2021.708178

Ostrovski, K. R. (2019). Desenvolvimento, produção e qualidade do mel de abelha Mandaçaia MQQ em ambientes urbano e rural. Universidade Federal do Paraná.

Peck, D. T., & Seeley, T. D. (2019). Mite bombs or robber lures? The roles of drifting and robbing in Varroa destructor transmission from collapsing honey bee colonies to their neighbors. PLoS ONE, 14(6), 1–14. doi: 10.1371/journal.pone.0218392

Pereira, F. M., Souza, B., & Lopes, M. T. (2017). Sistema de criação de abelhas-sem-ferrão (pp. 1–32). Embrapa Meio-Norte. Retrieved from https://www.infoteca.cnptia.embrapa.br/infoteca/bitstream/doc/1079116/1/CriacaoAbelhaSemFerrao.pdf%0A

Pfeiffer, K. J., & Crailsheim, K. (1998). Drifting of honeybees. Insectes Sociaux, 45, 151–167. doi: 020-1812/98/020151-17 $ 1.50+0.20/0

Purkiss, T., & Lach, L. (2019). Pathogen spillover from Apis mellifera to a stingless bee. Proceedings of the Royal Society B: Biological Sciences, 286(1908). doi: 10.1098/rspb.2019.1071

Quezada-Euán, J. J. G. (2018). Stingless Bees of Mexico. Springer Nature.

Retschnig, G., Kellermann, L. A., Mehmann, M. M., Yañez, O., Winiger, P., Williams, G. R., & Neumann, P. (2019). Black queen cell virus and drifting of honey bee workers (Apis mellifera). Journal of Apicultural Research, 58(5), 754–755. doi: 10.1080/00218839.2019.1655133

Roubik, D. W. (1989). Ecology and Natural History of Tropical Bees. Cambridge University Press.

Roubik, D. W. (2018). The pollination of cultivated plants: A compendium for practitioners. Food and Agriculture Organization of the United Nations.

Seeley, T. D. (2007). Honey bees of the Arnot Forest: a population of feral colonies persisting with Varroa destructor in the northeastern United States. Apidologie, 38(1), 19–29. doi: 10.1051/apido:2006055

Seeley, T. D., & Smith, M. L. (2015). Crowding honeybee colonies in apiaries can increase their vulnerability to the deadly ectoparasite Varroa destructor. Apidologie, 46(6), 716–727. doi: 10.1007/s13592-015-0361-2

Šekulja, D., Hermann, P., & Licek, E. (2014). Drifting behaviour of honeybees (Apis mellifera carnica Pollman, 1879) in the epidemiology of American fouldbrood. Zbornik Veleučilišta u Rijeci., 2(1), 345–358.

Silva-Neto, C. de M. e, Ribeiro, A. C. C., Gomes, F. L., Melo, A. P. C. de, Oliveira, G. M. de, Faquinello, P., Franceschinelli, E. V., & Nascimento, A. dos R. (2019). The stingless bee mandaçaia (Melipona quadrifasciata Lepeletier) increases the quality of greenhouse tomatoes. Journal of Apicultural Research, 58(1), 9–15. doi: 10.1080/00218839.2018.1494913

Silva, M., & Ramalho, M. (2016). The influence of habitat and species attributes on the density and nest spacing of a stingless bee (Meliponini) in the Atlantic Rainforest. Sociobiology, 63(3), 991–997. doi: 10.13102/sociobiology.v63i3.1037

Silva, P. I., Oliveira, F. A. S., Pedroza, H. P., Gadelha, I. C. N., Melo, M. M., & Soto-Blanco, B. (2015). Pesticide exposure of honeybees (Apis mellifera) pollinating melon crops. Apidologie, 46(6), 703–715. doi: 10.1007/s13592-015-0360-3

Slaa, E. J., Sánchez Chaves, L. A., Malagodi-Braga, K. S., & Hofstede, F. E. (2006). Stingless bees in applied pollination: practice and perspectives. Apidologie, 37(2), 293–315. doi: 10.1051/apido:2006022

Stephens, R. E., Beekman, M., & Gloag, R. (2017). The upside of recognition error? Artificially aggregated colonies of the stingless bee Tetragonula carbonaria tolerate high rates of worker drift. Biological Journal of the Linnean Society, 121(2), 258–266. doi: 10.1093/BIOLINNEAN/BLW048

Teixeira, É. W., Viana, T. A., Lima, M. A. P., Martins, G. F., & Lourenço, A. P. (2025). Detection and identification of Melissococcus plutonius in stingless bees (Apidae: Meliponini) from Brazil. Journal of Invertebrate Pathology, 213(July). doi: 10.1016/j.jip.2025.108418

Viana, B. F., Gabriel, J., Garibaldi, L. A., Gastagnino, L. B., Gramacho, K. P., & Oliveira, F. (2014). Stingless bees further improve apple pollination and production. Journal of Pollination Ecology, 14(25), 261–269.

Wei, C. A., & Dyer, F. C. (2009). Investing in learning: why do honeybees, Apis mellifera, vary the durations of learning flights? Animal Behaviour, 77(5), 1165–1177. doi: 10.1016/j.anbehav.2008.12.031

Wilms, W., Imperatriz-Fonseca, V. L., & Engels, W. (1996). Resource Partitioning between Highly Eusocial Bees and Possible Impact of the Introduced Africanized Honey Bee on Native Stingless Bees in the Brazilian Atlantic Rainforest. Studies on Neotropical Fauna and Environment, 31(3–4), 137–151. doi: 10.1076/snfe.31.3.137.13336

Zeil, J., Kelber, A., & Voss, R. (1996). Structure and function of learning flights in bees and wasps. Journal of Experimental Biology, 199(1), 245–252.

